# Measurement of the Persistence Length of Cytoskeletal Filaments using Curvature Distributions

**DOI:** 10.1101/252551

**Authors:** Pattipong Wisanpitayakorn, Keith J. Mickolajczyk, William O. Hancock, Luis Vidali, Erkan Tüzel

## Abstract

Cytoskeletal filaments such as microtubules and actin filaments play important roles in the mechanical integrity of cells and the ability of cells to respond to their environment. Measuring the mechanical properties of cytoskeletal structures is crucial for gaining insight into intracellular mechanical stresses and their role in regulating cellular processes. One of the ways to characterize these mechanical properties is by measuring their persistence length, the average length over which filaments stay straight. There are several approaches in the literature for measuring filament deformations, including Fourier analysis of images obtained using fluorescence microscopy. Here, we show how curvature distributions can be used as an alternative tool to quantify bio-filament deformations, and investigate how the apparent stiffness of filaments depends on the resolution and noise of the imaging system. We present analytical calculations of the scaling curvature distributions as a function of filament discretization, and test our predictions by comparing Monte Carlo simulations to results from existing techniques. We also apply our approach to microtubules and actin filaments obtained from *in vitro* gliding assay experiments with high densities of non-functional motors, and calculate the persistence length of these filaments. The presented curvature analysis is significantly more accurate compared to existing approaches for small data sets, and can be readily applied to both *in vitro* or *in vivo* filament data through the use of an ImageJ plugin we provide.

## 1. INTRODUCTION

The cytoskeleton is a filamentous network found in cells, which maintains cell shape, aids in cell motion, and plays a key role in intracellular transport and cell division (1); it is made up of microtubules, actin filaments, and intermediate filaments. Microtubules are the rigid tubular structures in cells that in addition to their structural role act like molecular highways to transport cellular signals and cargo in a wide variety of cellular processes (2). While microtubules are hair-like in terms of their aspect ratio, they have a large resistance to bending–same as Plexiglass (3). Actin filaments on the other hand are significantly softer and easier to bend compared to microtubules. They also serve a variety of roles in cells, including formation of protrusions at the leading edge of a crawling fibroblasts and contraction of the cytokinetic machinery (1). Utilizing cytoskeletal associated proteins and molecular motors, both microtubules and actin filaments can also form active or passive bundles such as flagella of swimming organisms or microvilli on the brush-border cells lining the intestine (1). Thus, it is important to study the mechanical behavior of these filaments to understand their function as dynamic mechanical components in living cells, and gain insight into the intracellular mechanical stresses (4).

Inside a cell, microtubules and actin filaments continuously bend and buckle, resisting forces as they perform their function; therefore, a filament’s resistance to bending is an important mechanical property to measure. One way to characterize the bending deformations of these filaments is to measure the persistence length, *L_p_*, the length scale of which thermal fluctuations can cause the filaments to spontaneously bend (5). One of the first estimations of microtubule persistence length *in vitro* made use of statistical analysis of the contour lengths and end-to-end distances in dark-field microscopic images, which resulted in a very small values of about 75*μ*m (6). The measured microtubule persistence length was an order of magnitude greater than that of actin filaments; nevertheless, it is much lower than the accepted values for microtubules of 1-10 *mm* in recent literature (7). Since then, microtubule and actin flexural rigidity (therefore persistence length) has been further estimated from dynamic video images using various techniques, including spectral analysis of thermally fluctuating filaments (3, 8–11), end fluctuations (12, 13), and tangent correlations (11, 14, 15). There have also been non-equilibrium measurements, including hydrodynamic flow (8, 12, 16), optical trapping (17–21), and atomic force microscopy (22).

Despite numerous studies on persistence length of bio-filaments *in vitro* in the past three decades (3, 6, 14, 15, 21, 23), far fewer measurements exist *in vivo*. Some of the existing approaches are not readily applicable *in vivo* due to the complexity of forces and the boundary conditions involved (24). The crowded environment of the cell also limits the number of filaments that can be traced in a given microscopy image. It is also a challenge to obtain long temporal resolution from an *in vivo* experiment to achieve accurate persistence length measurement (25). In addition, in living cells, these filaments can potentially have defects and associated proteins that can change local resistance to bending (26), complicating the interpretation of global methods such as Fourier analysis (3). It is, therefore, crucial to develop new approaches that focus on local deformations to measure the persistence length *L_p_* accurately, particularly applicable to small data sets.

To overcome some of these limitations, Odde et al. (27) used curvature distributions to describe the behavior of microtubules in the lamella of fibroblasts, and characterized the breaking mechanisms of microtubules. Since then, curvature distributions have been used to characterize bio-polymers, for example, to analyze DNA flexibility (28), to quantify the bias in branching direction of actin filaments (29), and to study microtubule bending under perpendicular electric forces (30). Bicek et al. (31) used curvature distributions of microtubules to study the anterograde transport of microtubules in LLC-PK1 epithelial cells, where exponential distributions of curvatures were observed *in vivo* when microtubules were driven by motor forces. Similar exponential distributions were observed in the analysis of actively driven actin filaments (32). However, obtaining quantitative results of the persistence length remains a challenge, given the effects of spatial sampling and experimental noise on the observed curvature distributions (4).

In this work we show how these challenges can be overcome by using a sub-sampling approach, combined with analytical calculations of the scaling of form for curvature distributions. We validate our approach using simulated, model-convolved (4, 33) filaments and compare it to the popular approach, the Fourier analysis (3). We apply our curvature analysis to measure the persistence lengths of microtubule and actin filaments immobilized on glass surfaces with non-functional motors. Finally, we illustrate how our approach is significantly more accurate than Fourier analysis when applied to typical *in vitro* and *in vivo* scenarios with varying length filaments and small data sets (< 20 filaments).

## 2. MATERIALS AND METHODS

### 2.1 Simulations oflaments

In order to compare the performance of the curvature analysis to the other existing techniques, 10 *μ*m long filaments with bond spacing of 100 *nm* were generated using Monte Carlo sampling of the Gaussian curvature distribution (as discussed in Section 3). Filaments, however, are often traced from experimental images, and the Point Spread Function (PSF) of light, together with contamination from detector noise and optical aberrations, make accurate collection of *x-y* coordinates difficult. To better compare with experimental data, we convolved the coordinates of simulated filaments with the PSF, and added experimental noise-an approach called Model Convolution Microscopy (4, 33, 34). More specifically, for each filament, the simulated coordinates were projected onto 10 *nm* two dimensional grid. We then used Bresenham’s line algorithm (35) to fill in the grid along those coordinates to create a pixellated polymer with radius corresponding to that of an actin filaments or microtubule. The polymer data was then convolved with a Gaussian PSF with a wavelength of 515 *nm* and numerical aperture (NA) of 1.4, in order to make every grid element along the backbone a point source of light. The noise from image acquisition with a digital camera was distributed randomly with mean and standard deviation measured from a typical experimental image obtained using our microscope systems. The fine grid was coarse grained to a larger grid size of 100 *nm*, approximately corresponding to the typical pixel size from a fluorescence microscope. This procedure creates an image of a simulated filament that is directly comparable to an experimental image (see **Fig. S1** in the **Supporting Material**).

### 2.2 Linear interpolation

In the simulations, the coordinates of each filament were considered to be equally spaced. However, the coordinate spacing of the filaments obtained from the tracing programs can be slightly non-uniform. Therefore, we perform linear interpolation adapted from Ott et al. (14) on the traced filaments to convert them into filaments with uniform coordinate spacing. The linear interpolation is started by drawing a circle of radius 100 *nm* around the first coordinates of the filament. The point where the circle intersects with the line connecting between two adjacent coordinates were recorded as new coordinates, which were used as the center of a new circle. The process was repeated until the end of the filament was reached. This linear interpolation procedure was performed on the filaments before undergoing curvature and Fourier analyses. Even though the process showed no significant effect on the Fourier mode variances, it slightly improved the curvature analysis as shown in **Fig. S2** in the **Supporting Material**.

### 2.3 Preparation and imaging of actin filaments

Chicken skeletal muscle actin was purified from acetonic powder using standard methods developed to purify actin from skeletal muscle (36) and gel filtered according to (37). Fractions containing actin were rapidly frozen in liquid nitrogen as 50 ml pellets and stored at -80°C. Myosin S1 fragment was prepared from chicken skeletal muscle (38) and the S1 preparation was also stored as 50 ml frozen pellets at -80°C. To visualize F-actin, actin filaments polymerized with 100 mM KCl and 5 mM MgCl_2_ from a 2 mM solution of monomers were decorated with a molar excess of Alexa-488-phalloidin (Molecular Probes) overnight, and diluted before use to 20 nM with AB buffer (25 mM imidazole-HCl, 25 mM KCl, 4 mM MgCl_2_, 1 mM EGTA, 1 mM DTT, pH = 7.4) plus 100 mM DTT (39). Coverslips were affixed to a slide by water droplets and coated with 0.2% (w/v) nitrocellulose in isoamyl acetate, and allowed to dry. Fifty to 100 *μ*L of S1 solution diluted in AB buffer at a various concentrations was pipetted onto parafilm on top of ice. The coverslip was placed with the nitrocellulose facing down onto the sample and incubated for an hour. The coverslip was then washed with 100 *μ*L of AB buffer supplemented with 0.5 mg/ml BSA to block any unoccupied nitrocellulose. A drop of 20 nM phalloidin-decorated actin was added to a slide and the coverslip containing the bound S1 was placed on the drop on the slide. Imaging was performed using a Zeiss Axiovert 200M microscope equipped with a CoolSNAP fx CDD camera. A filter cube for FITC/GFP was used with a 63X objective DIC Plan-APO CHROMAT NA 1.4. The microscope and camera were driven by Zeiss Axiovision software.

### 2.4 Preparation and imaging of microtubules

Cy-5-labeled microtubules were polymerized from bovine brain tubulin as previously reported (40) and stabilized with 10*μ*M taxol. A gliding assay was prepared using 500 pM rigor (R210A) full-length Drosophila melanogaster kinesin-1 heavy chain. 0.5 mg/mL casein was used both for blocking and for binding the rigor kinesin to the glass cover slip, as previously reported (41). The final imaging solution was 0.5 mg/mL casein, 10 *μ*M taxol, 20 mM glucose, 20*μ*g/mL glucose oxidase, 8*μ*g/mL catalase, 1:200 *β*-mercaptoethanol, and 2 mM ATP in BRB80 (80 mM PIPES, 1 mM EGTA, 1 mM MgCl2, pH 6.8). Experiments were performed under total internal reflection fluorescence using a Nikon TE2000 inverted microscope and a Melles Griot 632 nm Helium-Neon laser. Static images were captured using a Cascade 512 EMCCD camera (Roper Scientific) and MetaVue software (Molecular Devices) using a 400 ms exposure time. The pixel size was 108.1 nm/pix.

### 2.5 Filament tracing

The model-convolved and experimental filaments were traced with two types of tracing algorithms: an in-house developed Gaussian Scan (GS) approach and the ImageJ plug-in JFilament (11). GS was implemented in MATLAB closely following the approach by Brangwynne et al.(10). Briefly, the images are processed through noise reduction (median filter), followed by skeletonization. The filament backbones are then fit to a 4*^th^* order polynomial, which is then used to calculate the perpendicular angle along the backbone of the filament, and a scan along this axis is performed on the original image to obtain an intensity profile. Finally, these profiles were fit to a Gaussian function and the mean values of the functions were taken as the coordinates of the filaments. In contrast, JFilament is based on stretching open active contours or “snakes”—parametric curves that deform to minimize the sum of an external energy derived from the image and an internal bending and stretching energy (11). Two types of forces are introduced from the external energy: (i) forces that pull the contour toward the central bright line of a filament, and (ii) forces that stretch the contour toward the ends of bright ridges. To trace a filament, initial snake is given by hand-clicking along the filament, and it is then deformed to match the filament shape on the image. For further details, the reader is referred to Ref. (11).

### 2.6 Fourier analysis

The persistence length, *L_p_*, of a biopolymer can be calculated using a Fourier analysis approach introduced by Gittes et al. (3). In this approach, we can express the filament shape, characterized by *Φ(s)*, as a superposition of Fourier modes, <u>i.e.</u>,

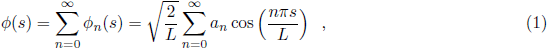

where *L* is the total length of the filament, *s* is the arc length along the filament, *n* is the mode number, and *a_n_* is the mode amplitude. For a thermally-driven polymer, the variance of the model amplitudes scales as ~*n*^−2^, however, experimentally obtained data introduces pixellation errors as position along a filament deviates from the actual position by a random distance *ϵ_k_*. Gittes et al. (3) showed that after taking pixellation into account, the variance of the mode amplitudes becomes

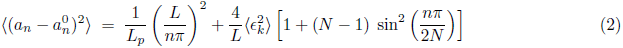

This modified variance formula is often used in practice to calculate L_p_. For data sets with varying filament lengths, however, this approach would not be adequate as it assumes a filament of given length *L* is measured as a function of time, and used in the Fourier analysis. In order to make a fair comparison with the curvature approach that we apply to filaments with different lengths, one can calculate a modified length-dependent variance that takes into account length variation

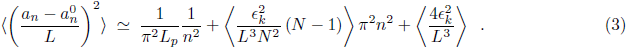

Note that this expression is valid for small *n* values. Nonetheless, since experimental noise is dominant at high *n* values, this approximation is valid for most practical situations.

### 2.7 Bootstrapping

To measure the persistence length accurately for small data sets, we adapted the bootstrapping approach proposed by Hawkins et al. (23). In our bootstrapping, the same number of filaments as the size of the set of filaments were randomly picked out of a given set (picking the same filament again is allowed). Those selected filaments then underwent curvature and Fourier analyses to estimate the persistence length. This process was repeated 5,000 times to construct a histogram of persistence lengths, and the mean persistence length and the associated standard deviation were estimated by fitting the histogram to a log-normal distribution.

## 3. CURVATURE DISTRIBUTION OF A FILAMENT

In order to use curvature distributions to measure *L_p_* quantitatively, some form of sub-sampling is required to avoid measurement noise, either due to pixellation or tracing algorithm induced errors (4). In what follows, starting with the Hamiltonian of the wormlike chain model, we analytically calculate how the curvature distribution rescales as a decimation procedure (as a way to sub-sample) is applied on the traced bio-polymer coordinates. We then obtain a theoretical expression for the variance of the curvature distribution that can be used to calculate the persistence length *L_p_* quantitatively. Validations of theoretical results with simulations, and application of this approach to experimental data is given in the Results section.

### 3.1 Wormlike chain model and curvature distributions

In the continuum description, a semi-flexible polymer such as an actin filament or microtubule (as illustrated in Fig. 1a) can be modeled using the wormlike chain model, with the Hamiltonian given by

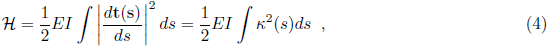

where *EI* is the flexural rigidity (*E*: Young’s modulus, *I*: second moment of cross sectional area), t_(s)_ is the unit tangent vector at *s*, and the curvature is defined as *κ ≡|dt(s)/ds|=dθ/ds*. Typically, a discretized form of this energy functional is used in either interpretation of experiments or in simulations, which can be written as

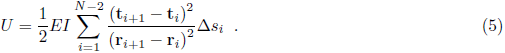

Note that the presence of the bond lengths in the denominator is crucial for proper discretization (42). The local curvature along the backbone of the chain can be simply calculated as

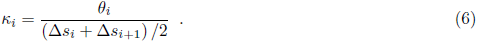

Assuming that bending energy is the dominant energy and using equi-partition one can show that curvatures are going to be distributed according to

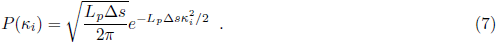

Here, the persistence length *L_p_* is related to the flexural rigidity via *L_p_ = EI/(k_B_T)*, were *k_B_* is Boltzmann’s constant, and *T* is the temperature (2). Similar to curvatures, the distribution of consecutive angles obeys a Gaussian distribution, i.e.

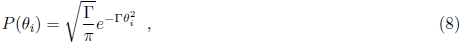

where Γ ≡ *L_p_*Δ*s*/2. While for ideal thermally-driven polymers the curvature distribution is given by Eq. 8, due to limits in resolution and inherent noise, and errors introduced by tracing algorithms, the curvature distribution for experimentally observed filaments often deviates from a Gaussian distribution. In order to eliminate contributions from noise, it has been proposed that some form of sub-sampling should be performed to analyze the curvature distribution outside of the noise-dominated regime (4). For the semi-flexible polymer model considered here, one way to accomplish this is to coarse-grain the polymer chain via a *decimation* procedure. The polymer after decimation has fewer nodes, and the resulting distribution of angles has to be calculated again accordingly. In what follows, we will show how the curvature distribution scales as a function of such decimation, first for the special case of *N* = 5 nodes, and later more generally to derive a scaling relationship.

### 3.2 Re-normalized curvature distribution for *N*= 5

A sample bio-polymer of *N* = 5 nodes is shown in *Fig. S3a* in the *Supporting Material*. After one step of decimation, i.e. removing every other node, there is only one bond angle left, namely 
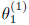
 The subscript on the angles denote the sequential numbering of the angles along the backbone, whereas the superscript shows the decimation level. For the bio-polymer shown in *Fig. S3a* only of step decimation is possible, which makes calculations easier. For simplicity we assume all the bond lengths in the original chain are equal and given by 
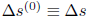
 and they scale as 
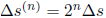
 where *n* shows the decimation level. This assumption implies that the bio-polymers considered here are fairly stiff, i.e. *L/Lp* ≪1, as in the case of microtubules. From Eq. 6 it follows that the local curvature at a node at a given decimation level *n* is given by

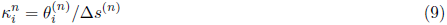

where *i* = 1,…, **M_n_*. Here M_n_ ≡ (N − 1)/2^n^* ∡ 1. For simplicity we will work with the bond angles instead of curvature in the following. Given the relatively stiff bio-polymer approximation, we consider the angle change between two consecutive bonds to be small. For the chain shown in **Fig. S3a**, in which the the first bond is aligned along the *x*-axis, using the small angle approximation one can easily show that

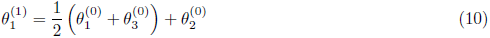

and therefore distribution of curvatures after one decimation can be written as

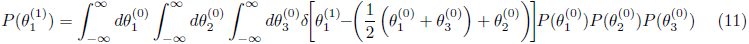

where 
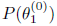
 is given by Eq. 8. Note that in principle the limits of integration depend on the particular configuration of the chain and the bond angles can only take values in the interval [−π/2; π/2]. Numerical evidence confirms, however, that the limits can be approximated by ±∞, even for relatively soft chains like actin filaments. Using the representation of the *δ*-function in Fourier space, one can show that

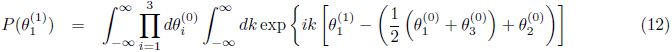

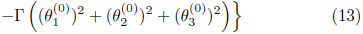

and evaluating the Gaussian integrals one gets

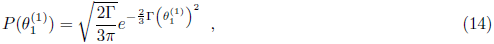

or in terms of the curvature

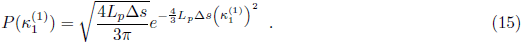

This result can be interpreted as rescaling of the persistence length 
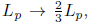
 and the bond length Δ*s* → 2^2^Δ*s*. It is also important to note that the angles chosen in **Fig. S3a** are all positive angles with respect to the bond projection, however, since the integration limits are from −∞ to ∞, the sign of the bond angle is irrelevant, i.e. Eq. 15 is valid independent of the coordinate system and sign of the angle change.

### 3.3 Generalization to arbitrary bio-polymer length and decimation level

Using the same arguments of the preceding section, it can be shown by induction that for a bio-polymer of *N* nodes (Fig. 1a), the bond angles at a decimation level *n* are related to the bond angles of the original chain, 
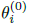
 as follows

**Figure 1.**
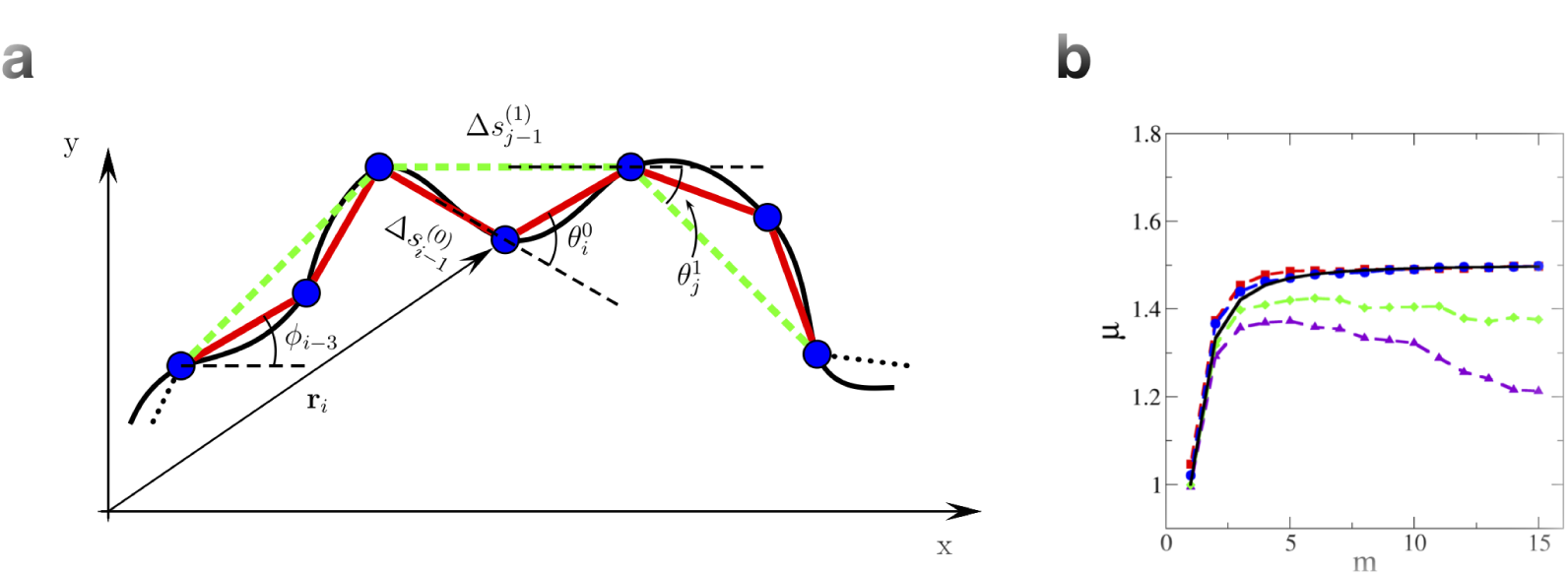
Illustration of the decimation process in a discrete polymer model, and validation of theoretical predictions using Monte Carlo simulations. (a) Continuum and discrete description of a bio-polymer. The distance from the origin to each node is denoted by r_*i*_, the angle between consecutive bonds is 
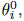
 and the distance between consecutive bonds is given by 
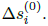
 Here, the superscript corresponds to the level of decimation performed. After one step decimation these bond angles and lengths are given by 
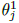
 and 
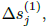
 respectively, and the dashed lines (green) show the new bio-polymer backbone after one step of decimation. (b) Plot of *μ*-the scaling factor of the persistence length due to sub-sampling-as a function of sub-sampling level, *m*, is shown for simulated filaments with different persistence length to pixel resolution ratios, *L_p_/Δs* = 100 (blue circles), 20 (red squares), 10 (green diamonds), and 5 (violet triangles), respectively. The black solid line shows the theoretical prediction, i.e. Eq. 33, where *m = 2^n^*.

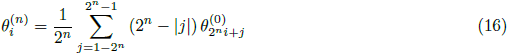

for the *i*th angle on the decimated bio-polymer, where *i* = 1,…, *M_n_*. Once again, since the integration limits are ±∞, the sign of the bond angles are irrelevant. For a bio-polymer of arbitrary length, as a result of the decimation procedure, the angles become correlated and the joint probability distribution of the bond angles at a decimation level *n* can be written as

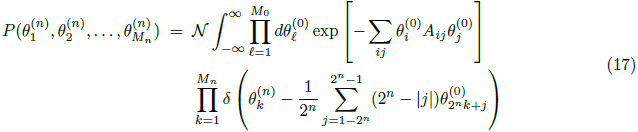

where *N* is the normalization constant. As before, the *δ*-functions in the above summation can be written in Fourier space resulting in *N* − 2 + *M_n_* dimensional integrals. However, the Gaussian integrals over the angle *θ*’s can be performed easily and using induction it can be shown that

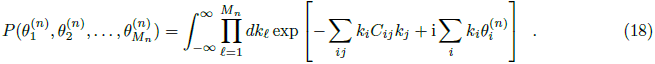

Here, C has the form

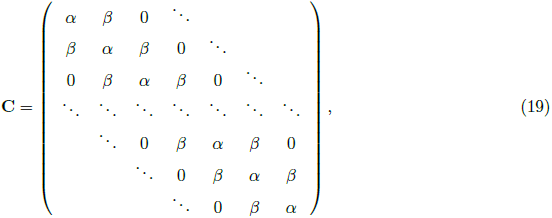

where

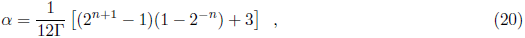

and

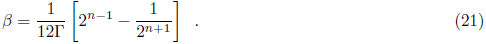

Performing the integrals in Eq. 18 one can show that the joint probability distribution is given by

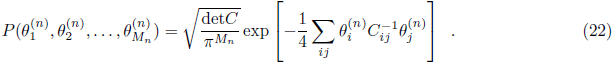

In principle Eq. 22 has all the necessary information about the bond angles and thereby curvatures on the underlying decimated bio-polymer. The tri-diagonal matrix C can be inverted using recursion relations (43). Alternatively, if a new matrix 
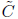
 is defined such that

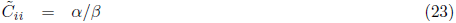

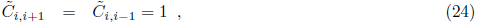

then it has been shown that the determinant and the matrix elements are given by (43)

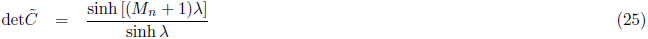

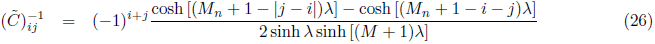

where 
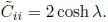

However, in a typical experiment, as we have discussed earlier, the curvatures along a given chain are measured and a distribution function is then calculated using these curvatures. This is equivalent to choosing a particular node, i.e. the first bond angle, and averaging the distribution of that bond angle with respect to the changes in all the other bond angles. This procedure can be formally written as an integral over all possible values the remaining bond angles can take, i.e.

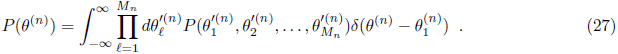

It can be easily shown that only *C*_11_ contributes and this integral reduces to 

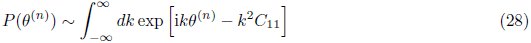

and using Eq. (20) one gets

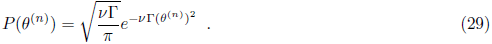

Here *ν* is given by

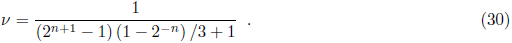

Writing the results in terms of curvature *κ^(n)^* one gets

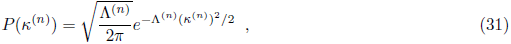

where the exponent Λ^(n)^ is

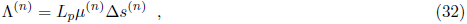

and *μ^(*n*)^* is found to be

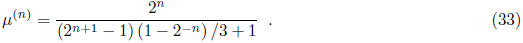

In the limit lim_n→∞_ *μ* = 3/2 which can be interpreted as a rescaled persistence length *L_p_* → (3/2)*L_p_*. This demonstrates that the effects on decimation can be significant especially if one is interested in quantitatively measuring the persistence length of the underlying chain. The slope of a plot of Λ^(*n*)^ (here Λ^(*n*)^ represents the inverse variance of the curvature distribution) as a function of *μ^(n)^* Δ*s^(n)^* (Eq. 32) can be used to calculate the value of the persistence length, *L_p_*, for a given semi-flexible polymer.

## 4. RESULTS

### 4.1 Theoretical predictions are recapitulated using simulations

In order to validate the theoretical results of the previous section, we generated bio-polymers with known persistence lengths using Monte Carlo simulations (see Materials and Methods). It is important to note that even though the calculations are done using a decimation procedure for simplicity, the results can be generalized to arbitrary sub-sampling where curvature distribution is measured using every *m^th^* node in a chain. In other words, Eqs. 31 to 33 that are in terms of *n*, are also true for the general case, where *n* = ln(*m*)/ ln(2) (i.e. *m* = 2*^n^*). The measured curvature distributions for 10 μm long filaments with a persistence length of 10 μm at two such sampling levels are shown in **Fig. S3b**. When calculated using every node (*m* = 1) and every 10^th^ node (*m* = 10), the observed distributions are both Gaussian as expected. This procedure of sub-sampling can be continued as long as the chains are long enough to measure a curvature value, and there is enough data to produce a histogram.

After fitting each of the curvature distributions from *m* = 1 to *m* = 15 to Eq. 31 for Λ^(*m*)^, *μ*^(*m*)^ was obtained by solving Eq. 32 using the input persistence length *L_p_*, and average bond spacing, Δ*s^(m)^* measured directly from the coordinates after sub-sampling. The measured μ^(*m*)^ values are in good agreement with the theoretical values calculated from Eq. 33, as shown in Fig. 1b, for a wide range of persistence lengths, despite the calculations being done in the limit of very stiff chains. In particular, our results show that the curvature approach performed very well when the persistence length is at least 20 times greater than the pixel resolution. For instance, if the images were taken using a fluorescence microscope with a pixel resolution of 100 *nm*, the presented approach would work well for filaments with a persistence length ≥ 2 *μ*m. Therefore, we expect the method to perform reasonably well for the two most commonly studied bio-polymers, namely, actin filaments and microtubules.

### 4.2 Curvature at short length scales is affected by experimental noise

To recapitulate a typical experimental measurement, we performed model convolution (4, 33) (see **Materials and Methods**) on the Monte Carlo generated filaments with input persistence lengths of 10 *μm* and 3 *mm*. These values were chosen to mimic actin- and microtubule-like filaments, respectively (3). The convolved filament images were then analyzed with two types of tracing algorithms: Gaussian Scan (GS) and JFilament (see **Materials and Methods**). The curvature distributions were measured recursively throughout the filaments for each sub-sampling level, *m*. As shown in Fig. 2b insets, the curvature distributions deviate from a Gaussian distribution for both soft (left panel) and stiff (right panel) GS traced filaments. Similar results were obtained when JFilament tracing is used for soft filaments (see **Fig. S4b** in the **Supporting Material**, left), showing that sub-sampling is required to avoid measuring curvature in the noise-dominated regime. It is important to note that while the distribution has a Gaussian form for JFilament traced of stiff filaments (**Fig. S4b, right**), the curvature distribution does not produce the theoretically expected variance. The curvature distributions of each sub-sampling level were then fitted to the Gaussian distribution in Eq. 31 to obtain Λ^(*m*)^. As observed in Fig. 2c, the curvature distributions for small sub-sampling levels were often affected by noise at low *μ^(m)^ Δs^(m)^* values, and deviate from a line. It is important note that here corresponds to the inverse variance of the curvature distribution at a given sub-sampling level, and therefore, higher values indicate narrowing of the curvature distribution.

**Figure 2.**
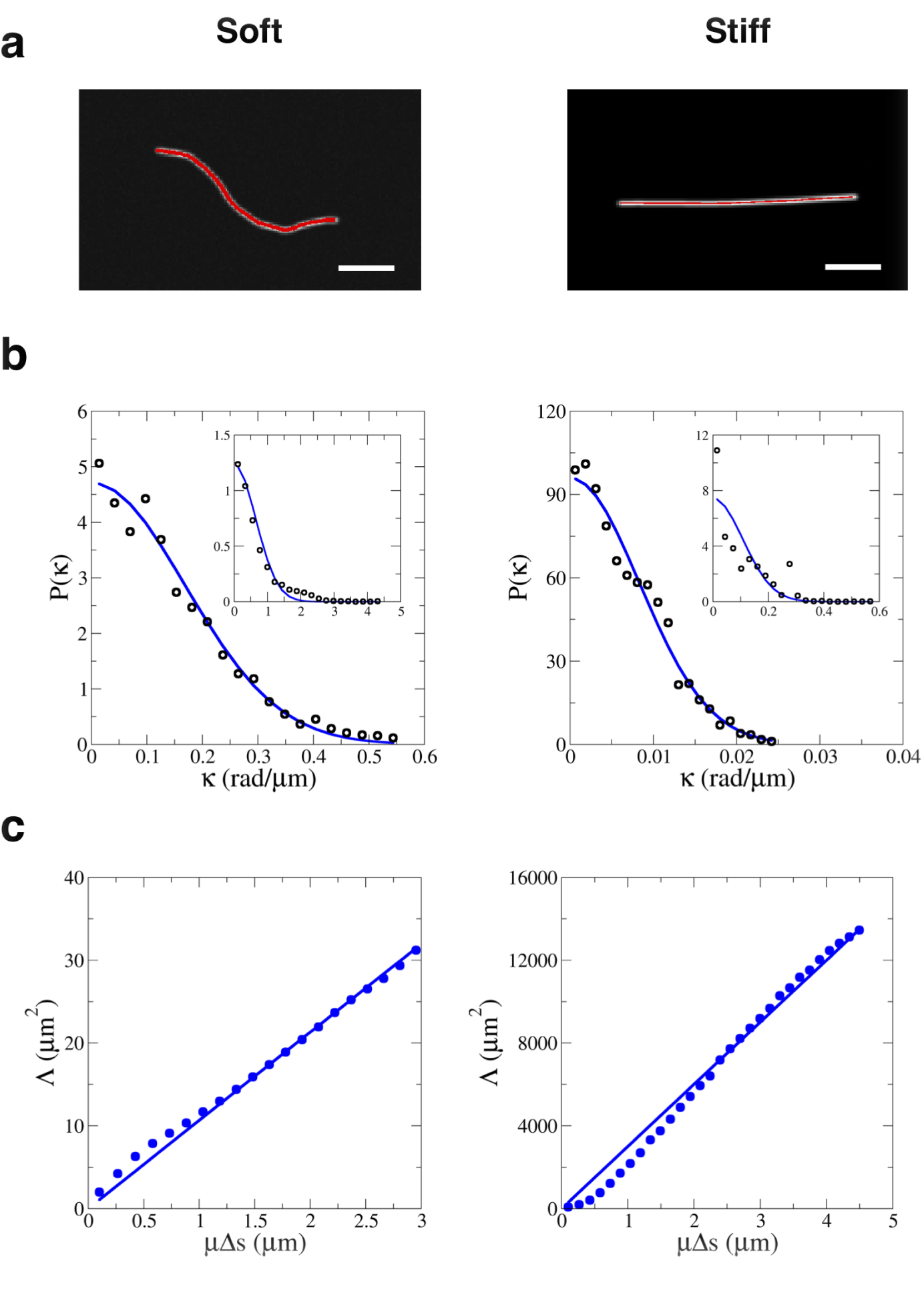
Validation of curvature analysis using model-convolved Monte Carlo simulated filaments with persistence lengths of 10 *μm* (left, soft) and 3 *mm* (right, stiff), traced by gaussian scan (GS) tracing. (a) Filament tracing. Red line represents recorded coordinates. (b) Curvature distribution of 100 filaments measured using every 20^th^ point along the backbone (*m* = 20) for the soft filaments, and every 30^th^ point along the backbone (*m* = 30) for the stiff filaments. The insets show the curvature distributions without sub-sampling (*m* = 1). Points denote simulation data, and the solid line is a fit to Eq. 31. (c) Plots of Λ^(*m*)^ as a function of *μ^(m)^ Δs^(m)^*, calculated using 100 filaments. Points represent data calculated using *m* = 1 to 20 (left panel), and *m* = 1 to 30 (right panel). Solid lines represent the fits of all data points to Eq. 32. Insets show GS tracing obtained contours (red) overlaid on the model-convolved filaments. The scale bar is 2 *Δm*.

The accuracy of curvature analysis was investigated over four different sampling sizes, namely, 10, 20, 50 and 100 filaments per set. For each sampling size, the curvature distribution for each sub-sampling level was measured for Λ^(*m*)^, and the measured values for Λ^(*m*)^ were fit to Eq. 32 to obtain *L_p_*. We also measured the persistence length using a modified Fourier analysis approach to allow for the analysis of filaments with varying lengths (see **Materials and Methods**). **Figure S5a** in the **Supporting Material** illustrates the variance plot of the Fourier mode amplitudes for simulated soft (left) and stiff (right) filaments. We also observed that the pixelation noise (concave-up tail) is more apparent on the variance plot for stiff filaments compared to the soft ones, as expected.

Since large numbers of filament coordinates are often difficult to obtain from experiments, the performance of the curvature and Fourier analyses were also tested using different filament set sizes, as shown in Fig. 3. The bar charts are each averaged over 10 independent sets of filaments, with error bars representing the standard deviations. The curvature analysis performed as accurately as the Fourier approach on large data sets as expected, but significantly better when analyzing smaller sets of filaments. Curvature distribution also provides more consistent results as indicated by smaller error bars. We observed no significant difference in results when performing theses analyses on the same filaments traced using JFilament (**Fig. S6b** in the **Supporting Material**).

**Figure 3.**
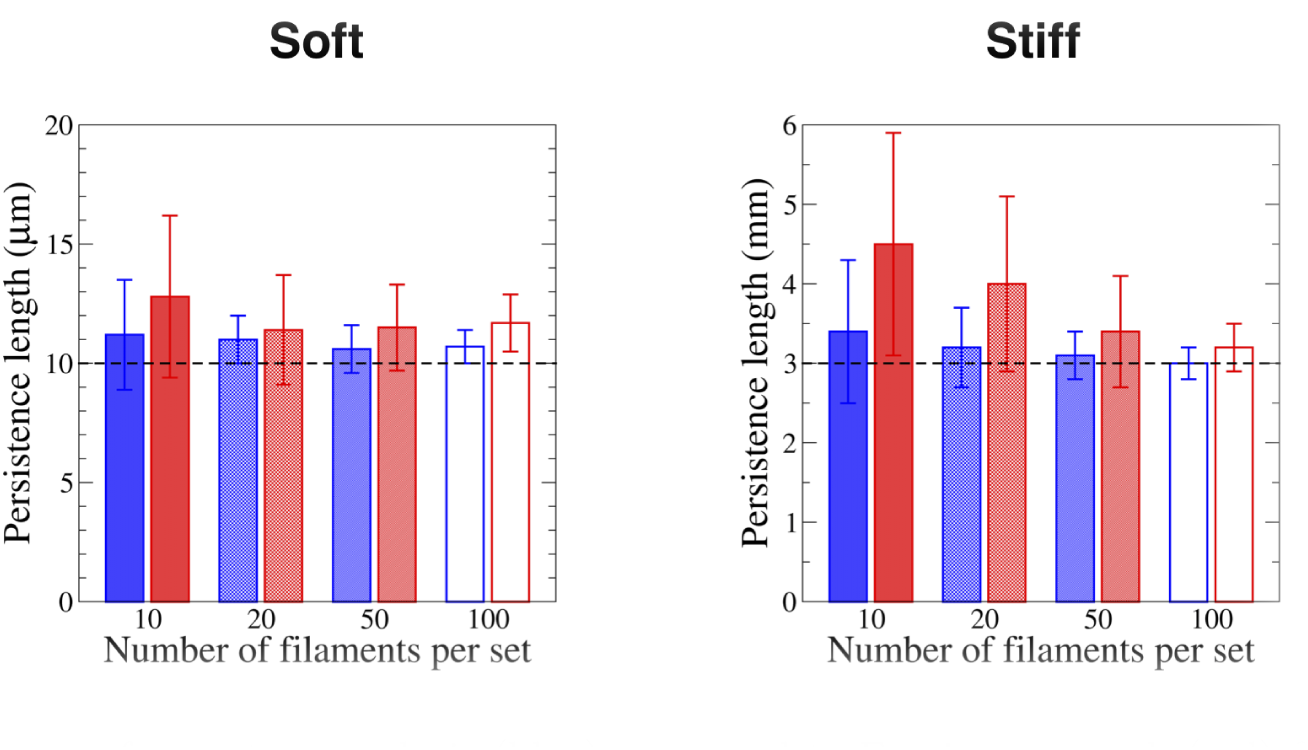
Accuracy of curvature analysis (blue) compared to Fourier approach (red) for different size data sets. All filaments are generated via Monte Carlo simulations, convolved with the Point Spread Function of the microscope, and traced by GS tracing. Each bar is an average over 10 sets of measurements with error bars representing standard deviation. The dashed lines represent the input persistence length.

### 4.3 Curvature distributions of experimental actin filaments and microtubules

The curvature analysis was then performed to measure the persistence length of actin filaments and microtubules *in vitro*, using filaments immobilized on glass surfaces with relatively high density of rigor (“non-functional”) motors (see **Materials and Methods**). effectively adhering the filaments to the surface allowed for image acquisition at slower frame rates, and helped avoid blurry images, in addition to preventing filament bundling over time. The filaments were traced using GS tracing and JFilament, as demonstrated in Fig. 4a and **Fig. S7a** in the **Supporting Material**. Since the results from our simulations suggested that both the curvature and Fourier analyses are at most accurate when analyzing large data sets, we implemented the bootstrapping method described by Hawkins et al. (23) (see **Materials and Methods**).

For each bootstrapped set, we measured the curvature distribution of those filaments at different sub-sampling levels and construct the Λ^(*m*)^ plot to find the persistence length. The distributions of measured *L_p_* values over 5, 000 bootstrapped sets are shown in Fig. 4c and **Fig. S7c**, where the histograms were fit to the log-normal distribution to calculate the mean and standard deviations. We found the persistence length of the actin filaments and microtubules traced via GS tracing to be 10.0 ± 0.7 *μm* and 300 ± 50 *μm*, respectively. The results for JFilament traced filaments were well within measurement error as shown in **Fig. S7c**. The filaments were also analyzed via Fourier analysis (**Fig. S8**), and the measured *L_p_* values obtained using both approaches were found to be comparable, for both types of filaments traced using GS (Fig. 5). While the *L_p_* value for actin filaments, traced with JFilament, was in agreement with the curvature analysis results (**Fig. S9**, also in agreement with GS tracing), the comparison for microtubules did not yield satisfactory results. Even though the curvature analysis gave similar results between two tracing approaches, the Fourier analysis performed poorly as evidenced by the variance plot in **Fig. S10a** in the **Supporting Material**, resulting in an underestimation of the persistence length.

**Figure 4.**
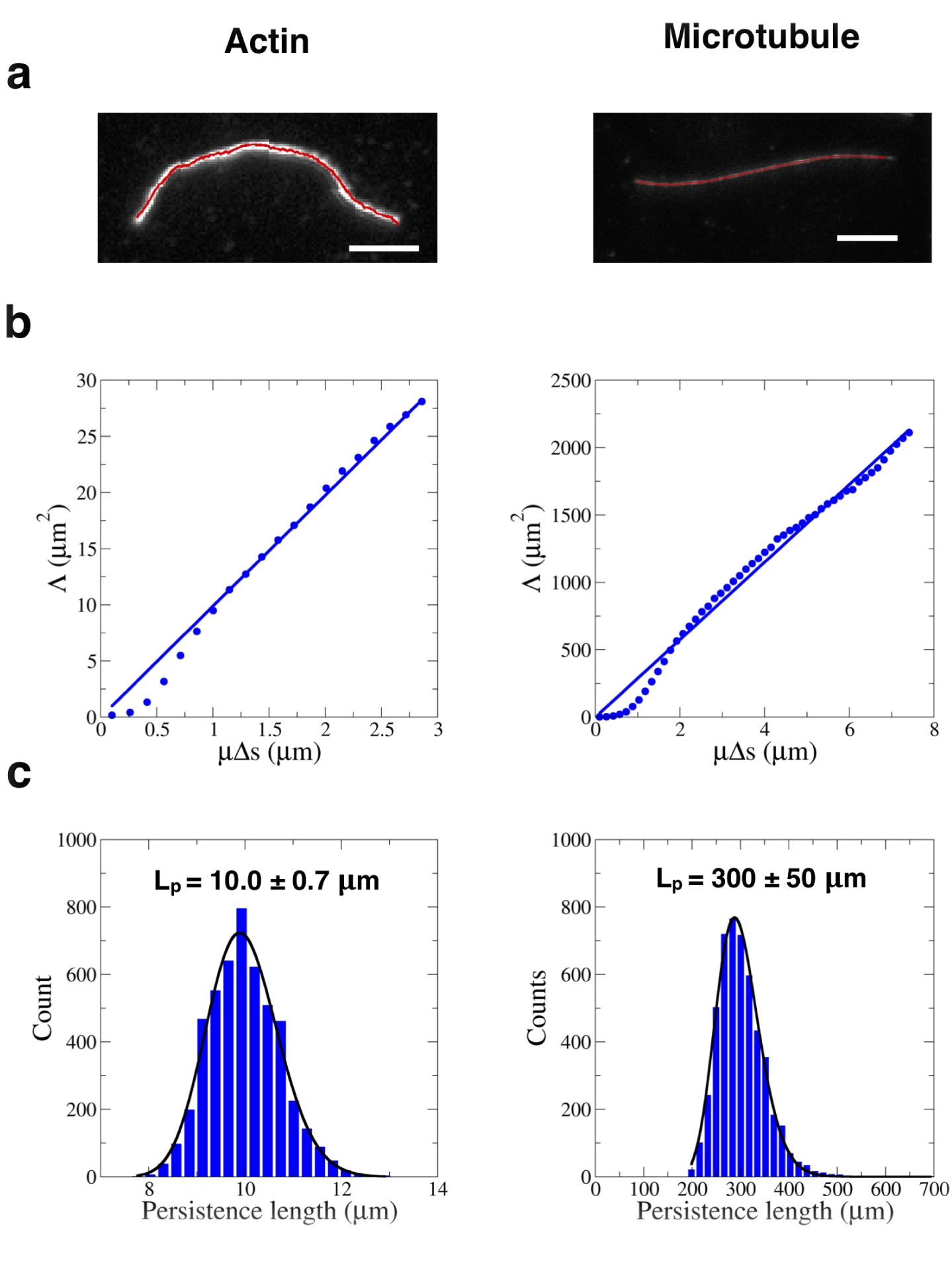
Curvature analysis applied to actin filaments (left) and microtubules (right) adhered to glass substrates with rigor/non-functional motors. All filament tracing is done via GS tracing. (a) Illustration of tracing of actin filaments and microtubules. Scale bar is 1 *μm* for the actin, and 2 *μm* for the micro-tubule. (b) Plots of Λ^(*m*)^ as a function of *μ^(m)^ Δs^(m)^*, calculated using 160 filaments. Blue dots represent data points calculated from actin filaments (left panel) using *m* = 1 to 20, and microtubules (right panel) using *m* = 1 to 50; blue lines represent fits to Eq. 32. (c) Histograms show distributions of the measured persistence length values obtained from bootstrapping 5, 000 data sets. The distributions were fit to the log-normal distribution to calculate the mean and standard deviation values.

**Figure 5.**
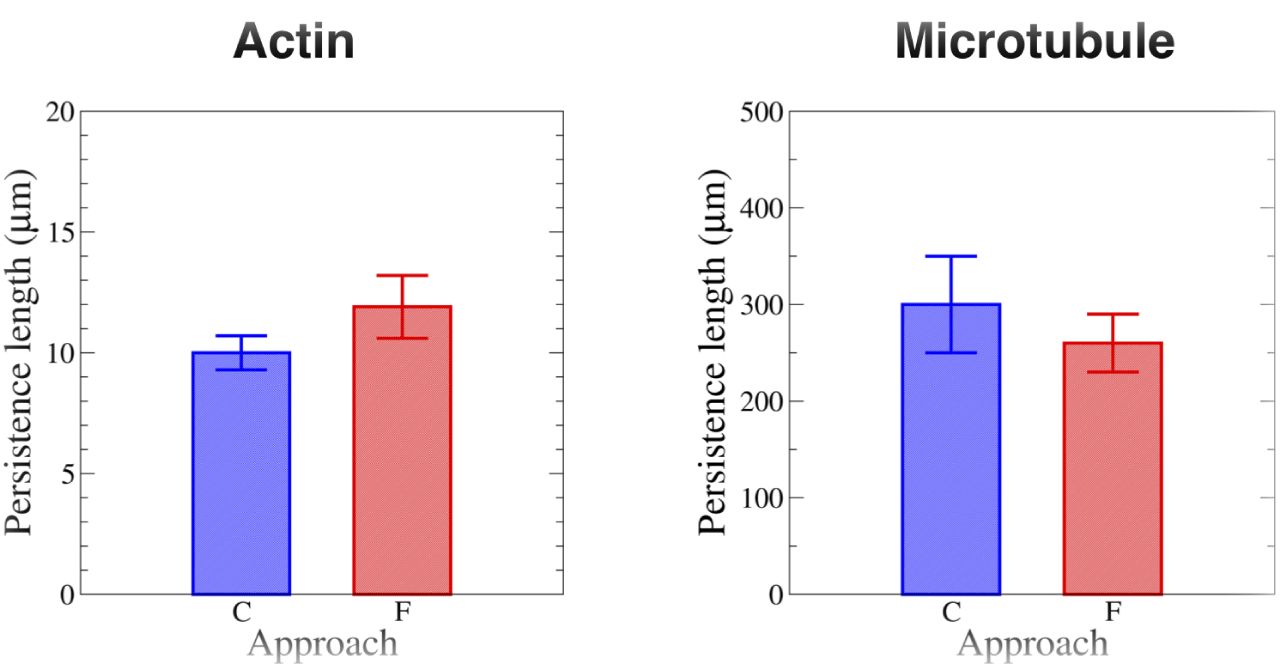
Comparison between persistence lengths measured with the curvature approach (marked C, blue) and Fourier analysis (marked F, red) for experimental actin filaments and microtubules, traced by GS scan.

### 4.4 Effect of length distribution and small data sets on persistence length measurement

It is often challenging to obtain a large number of filaments from fluorescence images *in vivo*, and in particular it is usually difficult to trace a given filament for a long enough time to acquire uncorrelated data for persistence length analysis. In addition, experimentally observed filaments often have length distributions, as shown in **Fig. S11** in the **Supporting Material**. To study the effect of length variation on persistence length analysis, we generated actin filaments and microtubules (*L_p_* = 10 *μm* and 300 *μm*, respectively) with experimentally observed length distributions *in vitro*, and used model convolution to generate realistic images (33). After tracing these filaments with GS tracing and JFilament, we applied curvature and our length-corrected Fourier analyses (see **Materials and Methods**). The measured persistence lengths from both analysis approaches and tracing methods are in excellent agreement with the input *L_p_* values for large number of filaments (> 50 filaments per analysis), suggesting that the curvature approach and our modified Fourier analysis can provide satisfactory results when applied to filaments with length variation (see **Fig. S12** and **Fig. S13** in the **Supporting Material**).

In order to investigate the effect of small number of filaments on analysis, we created 10 data sets, each containing 10 filaments generated and traced using the approaches described above. Insets of Fig. 6a and Fig. 6b show samples of such sets for actin- and microtubule-like filaments, respectively. As shown in Fig. 6a, Fig. 6b (and also in **Fig. S12c** and **Fig. S13c**), the curvature analysis clearly outperformed Fourier analysis under these conditions, making it more suitable for *in vivo* applications, especially when it is difficult to obtain large number of filaments for analysis.

**Figure 6.**
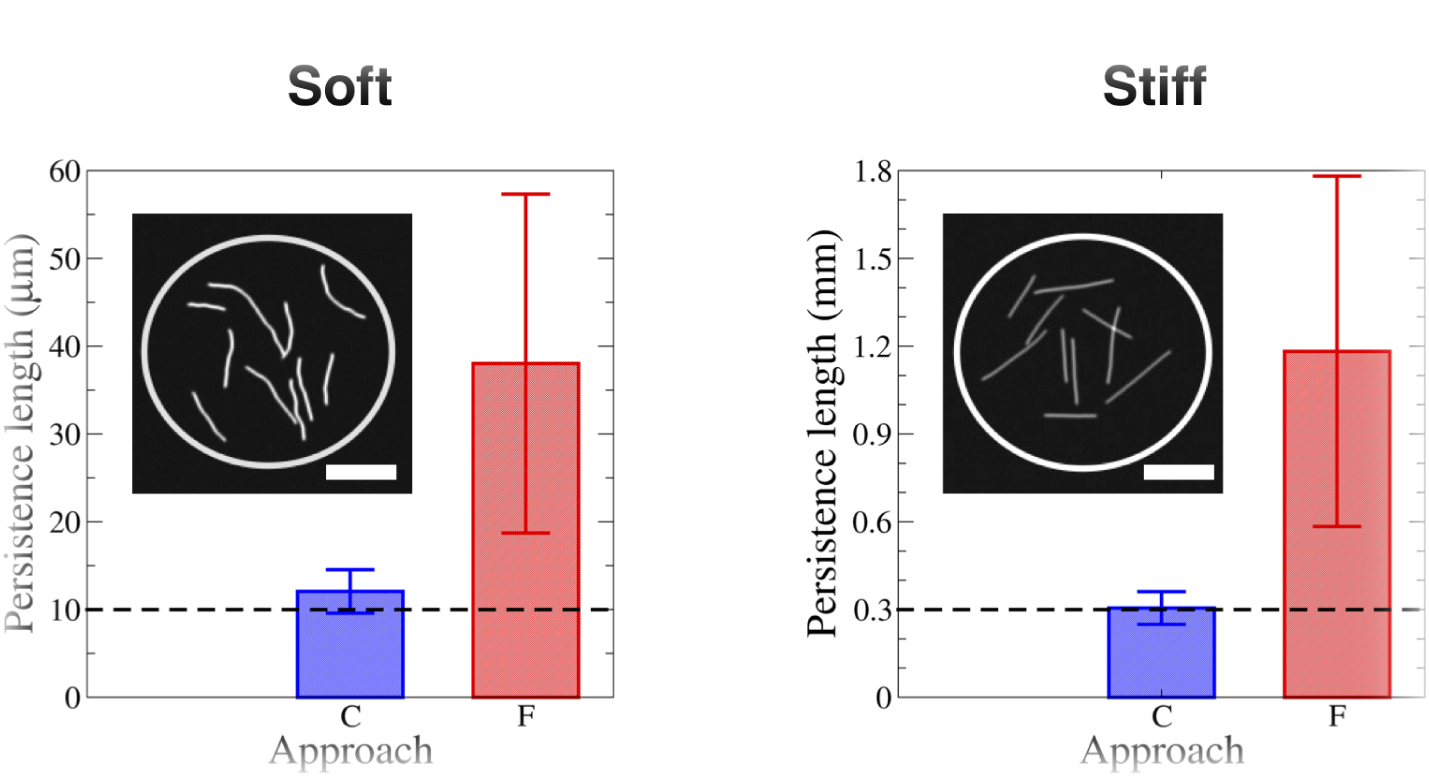
Curvature approach applied to small data sets of simulated actin filaments and microtubules of varying length, compared to length-corrected Fourier analysis. Ten sets each containing ten independent Monte Carlo generated filaments were used, and the contours were convolved with the Point Spread Function (PSF) of the microscope to create realistic images (samples shown in the insets). The filaments followed the same length distribution as the experimental filaments, but ones longer than 15 *μm* were discarded. Bars show the corresponding *L_p_* values measured using both approaches. The filaments were traced using the GS tracing. The scale bars are 5 *μm*.

## DISCUSSION

In this work, we developed a novel quantitative approach that relies on local curvature to measure the persistence length of cytoskeletal filaments. While in the past decade, we and others have used curvature analysis to characterize filament deformations (4, 31, 44), to the best of our knowledge, this is the first-time that curvature distributions have been used to quantify the persistence length of cytoskeletal filaments. More specifically, we analytically derived the scaling form of curvature distributions as a function of sub-sampling, a necessary step to avoid measurement noise. The accuracy of this theory was verified through model-convolved (33, 45) Monte-Carlo simulated actin- and microtubule-like filaments. We also investigated various factors such as tracing algorithms (Gaussian scan, JFilament (11)) and data set sizes. We compared our findings with the popular Fourier analysis approach (3, 9, 23, 25, 46) and found the results in excellent agreement.

When applied to an ensemble of filaments with varying filament lengths, we found that our curvature analysis was more accurate than existing approaches. It is important to note that in order to use the Fourier analysis (3, 23) with filaments of varying lengths, we had to modify the variance formulas in Ref. (3) to account for such length changes. Overall, we found that when there are sufficient filament coordinates, i.e. large data sets, both curvature and Fourier analysis approaches showed excellent agreement regardless of the tracking techniques used and the stiffness of the filaments. When our curvature analysis was applied to actin filaments and microtubules in *in vitro* gliding assays, the calculated persistence lengths were in good agreement with literature values (14, 18, 47–49). It is interesting to note that the rigidity of Taxol stabilized microtubules is about an order of magnitude less than what is commonly predicted for dynamic microtubules, supporting the hypothesis that Taxol softens microtubules (12, 16, 49).

Persistence length of a biological filament is an important quantitative descriptor of its resistance to bending, key to understanding the forces exerted on or by cytoskeletal filaments. While numerous approaches have been developed and used to measure persistence length of actin and microtubules in the past several decades (3, 6, 14, 21, 30, 50), there is considerable variability in reported values (4, 7) even from *in vitro* experiments. Many factors contribute to this discrepancy, including poor signal-to-noise ratios in fluorescence images giving rise to insufficient temporal resolution, variability in experimental conditions, limitations of measurement or filament tracking techniques, broad filament length distributions and limited data set sizes. Primarily due to these limitations, measurements of persistence length *in vivo* are scarce, particularly for stiff filaments such as microtubules (4).

The strength of the curvature analysis is its ability to provide significantly better accuracy for low number of filaments. This is particularly important for *in vivo* data, where obtaining large data sets under the same physiological conditions is often challenging (31, 46) or when investigating questions like the length dependent rigidity of microtubules(7, 50)-where binning filaments with respect to length reduces the number of data points significantly. It can also be used with electron microscopy images, for instance when determining the bending rigidity of individual microtubule protofilaments. While the example applications of the curvature analysis in this paper are limited to the two major cytoskeletal filaments, the approach can also be used to study bending deformations of any type polymers, such as DNA/RNA, flagella and cilia of motile organisms. In order to make this analysis more accessible, we developed an ImageJ plugin-Persistence Length Analyzer (PLA)-which is freely available at http://pla.tuzelgroup.net. This plugin allows the user to utilize both the curvature and Fourier analyses to measure the persistence length of given set of filament coordinates.

## AUTHOR CONTRIBUTIONS

PW developed the simulations and GS tracing software, performed the calculations and image tracing, wrote the ImageJ plugin, and the manuscript. LV and KJM performed the actin and microtubule experiments. WOH supervised the microtubule experiments. WOH, LV, ET contributed to the writing of the manuscript. ET conceived the study, wrote the manuscript and supervised the overall study.

## ACKNOWLEDGEMENTS

This work was supported by National Science Foundation grants CBET 1309933, and NSF-MCB 1253444, and National Institute of Health grants No. R01GM100076 and No. R01GM121679 to W.O.H and E.T.ET and LV also acknowledge support from Worcester Polytechnic Institute Startup Funds. We acknowledge the Thai government for supporting PW through the Development and Promotion of Science and Technology (DPST) Scholarship. The authors would also like to thank all members of the Tüzel, Vidali and Hancock Labs, Dan Sackett (NIH), Daniel M. Kroll (NDSU), David Odde (University of Minnesota) for helpful discussions and their insightful suggestions.

## References

1. Alberts, B., J. Alexander, J. Lewis, M. Raff, K. Roberts, and P. Walter, 2002. Molecular Biology of the Cell, Fourth Edition. Garland Science, New York.

2. Howard, J., 2001. Mechanics of Motor Proteins and the Cytoskeleton. Sinauer Associates, Inc., Sunderland, MA.

3. Gittes, F., B. Mickey, J. Nettleton, and J. Howard, 1993. Flexural rigidity of microtubules and actin filaments measured from thermal fluctuations in shape. J. Cell Biol 120:923–923.

4. Bicek, A. D., E. Tüzel, D. M. Kroll, and D. J. Odde, 2007. Analysis of microtubule curvature. Methods Cell Biol. 83:237–268.

5. Husson, J., L. Laan, and D. Dogterom, 2009. Force generation by microtubule bundles. Phys. Rev. Lett. 4:33–43.

6. Mizushima-Sugano, J., T. Maeda, and T. Miki-Noumura, 1983. Flexural rigidity of singlet microtubules estimated from statistical analysis of their contour lengths and end-to-end distances. BBA-Gen Subjects 755:257–262.

7. Hawkins, T., M. Mirigian, M. S. Yasar, and J. L. Ross, 2010. Mechanics of microtubules. Journal of biomechanics 43:23–30.

8. Kurz, J. C., and Jr R. C. Williams, 1995. Microtubule-associated proteins and the flexibility of microtubules. Biochemistry 34:13374–13380.

9. Mickey, B., and J. Howard, 1995. Rigidity of microtubules is increased by stabilizing agents. J. Cell Biol. 130:909–917.

10. Brangwynne, C. P., G. H. Koenderink, E. Barry, Z. Dogic, F. C. MacKintosh, and D. A. Weitz, 2007. Bending dynamics of uctuating biopolymers probed by automated high-resolution filament tracking. Biophys.J. 93:346–359.

11. Smith, M. B., H. Li, T. Shen, X. Huang, E. Yusuf, and D. Vavylonis, 2010. Segmentation and tracking of cytoskeletal filaments using open active contours. Cytoskeleton 67:693–705.

12. Venier, P., A. C. Maggs, M.-F. Carlier, and D. Pantaloni, 1994. Analysis of microtubule rigidity using hydrodynamic flow and thermal fluctuations. J. Biol. Chem 269:13353–13360.

13. Cassimeris, L., D. Gard, P. Tran, and H. P. Erickson, 2001. Xmap215 is a long thin molecule that does not increase microtubule stiffness. J. Cell Sci. 114:25–3033.

14. Ott, A., M. Magnasco, A. Simon, and A. Libchaber, 1993. Measurement of the persistence length of polymerized actin using fluorescence microscopy. Phys. Rev. E. 48:1642–1645.

15. Takatsuki, H. E. Bengtsson, and A. Månsson, 2014. Persistence length of fascin-cross-linked actin filament bundles in solution and the *in vitro* motility assay. BBA-Gen Subjects 1840:1933–1942.

16. Dye, R. B., S. P. Fink, and R. C. Williams, 1993. Taxol-induced flexibility of microtubules and its reversal by map-2 and tau. J. Biol. Chem. 268:6847–6850.

17. Kurachi, M., M. Hoshi, and H. Tashiro, 1995. Buckling of a single microtubule by optical trapping forces: direct measurement of microtubule rigidity. Cell Motil. Cytoskeleton 30:221–228.

18. Felgner, H., R. Frank, and M. Schliwa, 1996. Flexural rigidity of microtubules measured with the use of optical tweezers. J. Cell Sci. 109:509–516.

19. Felgner, H., R. Frank, J. Biernat, E.-M. Mandelkow, E. Mandelkow, B. Ludin, A. Matus, and M. Schliwa, 1997. Domains of neuronal microtubule-associated proteins and flexural rigidity of microtubules. J. Cell Biol. 138:1067–1075.

20. Takasone, T., S. Juodkazis, Y. Kawagishi, A. Yamaguchi, S. Matsuo, H. Sakakibara, H. Nakayama, and Misawa, 2002. Flexural rigidity of a single microtubule. Jpn. J. Appl. Phys..41:3015–3019.

21. Kikumoto, M., M. Kurachi, V. Tosa, and H. Tashiro, 2006. Flexural rigidity of individual microtubules measured by a buckling force with optical traps. Biophys. J. 90:1687–1696.

22. Kis, A., S. Kasas, B. Babić, A. Kulik, W. Benoit, G. Briggs, C. Schonenberger, S. Catsicas, and L. Forro, 2002. Nanomechanics of microtubules. Phys. Rev. Lett89:248101.

23. Hawkins, T. L., M. Mirigian, J. Li, M. S. Yasar, D. L. Sackett, D. Sept, and J. L. Ross, 2012. Perturbations in microtubule mechanics from tubulin preparation. Cell Mol. Bioeng 5:227–238.

24. Brangwynne, C. P., G. H. Koenderink, F. C. MacKintosh, and D. A. Weitz, 2008. Nonequilibrium microtubule fluctuations in a model cytoskeleton. Phys. Rev. Lett 100:118104.

25. Brangwynne, C. P., F. MacKintosh, and D. A. Weitz, 2007. Force fluctuations and polymerization dynamics of intracellular microtubules. Proc. Natl. Acad. Sci. USA 104:16128–16133.

26. Phillips, R., J. Kondev, J. Theriot, and H. Garcia, 2013. Physical Biology of the Cell. Garland Science, New York.

27. Odde, D. J., L. Ma, A. H. Briggs, A. DeMarco, and M. W. Kirschner, 1999. Microtubule bending and breaking in living fibroblast cells. J. Cell Sci. 112:3283–3288.

28. van Noort, J., T. van der Heijden, M. de Jager, C. Wyman, R. Kanaar, and C. Dekker, 2003. The coiled-coil of the human rad50 dna repair protein contains specific segments of increased flexibility. Proc. Natl. Acad. Sci. USA 100:7581–7586.

29. Risca, V. I., E. B. Wang, O. Chaudhuri, J. J. Chia, P. L. Geissler, and D. A. Fletcher, 2012. Actin filament curvature biases branching direction. Proc. Natl. Acad. Sci. USA 109:2913–2918.

30. Van den Heuvel, M., M. De Graaff, and C. Dekker, 2008. Microtubule curvatures under perpendicular electric forces reveal a low persistence length. Proc. Natl. Acad. Sci. USA 105:7941–7946.

31. Bicek, A. D., E. Tüzel, A. Demtchouk, M. Uppalapati, W. O. Hancock, D. M. Kroll, and D. J. Odde, 2009. Anterograde microtubule transport drives microtubule bending in llc-pk1 epithelial cells. Mol. Biol. Cell 20:2943–2953.

32. Weber, C. A., R. Suzuki, V. Schaller, I. S. Aranson, A. R. Bausch, and E. Frey, 2015. Random bursts determine dynamics of active filaments. Proc. Natl. Acad. Sci. USA 112:10703–10707.

33. Stottrup, B. L. A. H. Nguyen, and E. Tüzel, 2010. Taking another look with fluorescence microscopy: Image processing techniques in langmuir monolayers for the twenty-first century. BBA-Biomembranes 1798:1289–1300.

34. convolution: a computational approach to digital image interpretation, M., 2010. Model convolution: a computational approach to digital image interpretation. Cel. Mol. Bioeng. 3:163–170.

35. Bresenham, J. E., 1965. Algorithm for computer control of a digital plotter. IBM Syst. J 4:25–30.

36. Pardee, J. D., and J. A. Spudich, 1982. Purification of muscle actin. Methods Enzymol. 85:164–181.

37. MacLean-Fletcher, S., and T. D. Pollard, 1980. Identification of a factor in conventional muscle actin preparations which inhibits actin filament self-association. Biochem. Biophys. Res. Commun. 96:18–27.

38. Margossian, S. S., and S. Lowey, 1982. Preparation of myosin and its subfragments from rabbit skeletal muscle. Methods Enzymol. 85:55–71.

39. Kron, S. J., Y. Y. Toyoshima, T. Q. Uyeda, and J. A. Spudich, 1991. Assays for actin sliding movement over myosin-coated surfaces.Methods Enzymol. 196:399–416.

40. Shastry, S., and W. O. Hancock, 2010. Neck linker length determines the degree of processivity in kinesin-1 and kinesin-2 motors. Curr. Biol. 20:939–943.

41. Mickolajczyk, K. J., N. C. De enbaugh, J. O. Arroyo, J. Andrecka, P. Kukura, and W. O. Hancock, 2015. Kinetics of nucleotide-dependent structural transitions in the kinesin-1 hydrolysis cycle. Proc. Natl. Acad. Sci. USA 112:E7186–E7193.

42. Klapper, I., and H. Qian, 1998. Remarks on discrete and continuous large-scale models of dna dynamics. Biophys. J. 74:2504–2514.

43. Hu, G., and R. F. O’Connell, 1996. Analytical inversion of symmetric tridiagonal matrices. J. Phys. A: Math. Gen. 29:1511–1513.

44. Xu, T., D. Vavylonis, F.-C. Tsai, G. H. Koenderink, W. Nie, E. Yusuf, I.-J. Lee, J.-Q. Wu, and X. Huang, 2015. Soax: a software for quantification of 3d biopolymer networks. Sci. Rep. 5:9081.

45. Gardner, M. K., B. L. Sprague, C. G. Pearson, B. D. Cosgrove, A. D. Bicek, K. Bloom, E. Salmon, and D. J. Odde, 2010. Model convolution: a computational approach to digital image interpretation. Cell Mol. Bioeng 3:163–170.

46. Pallavicini, C., V. Levi, D. E. Wetzler, J. F. Angiolini, L. Bense~nor, M. A. Despósito, and L. Bruno, 2014. Lateral motion and bending of microtubules studied with a new single-filament tracking routine in living cells. Biophys. J. 106:2625–2635.

47. Isambert, H., P. Venier, A. C. Maggs, A. Fattoum, R. Kassab, D. Pantaloni, and M.-F. Carlier, 1995. Flexibility of actin filaments derived from thermal fluctuations. effect of bound nucleotide, phalloidin, and muscle regulatory proteins. J. Biol. Chem 270:11437–11444.

48. Van den Heuvel, M., S. Bolhuis, and C. Dekker, 2007. Persistence length measurements from stochastic single-microtubule trajectories. Nano Lett 7:3138–3144.

49. Kawaguchi, K., S. Ishiwata, and T. Yamashita, 2008. Temperature dependence of the flexural rigidity of single microtubules. Biochemical and biophysical research communications 366:637–642.

50. Pampaloni, F., G. Lattanzi, A. Jonáš, T. Surrey, E. Frey, and E.-L. Florin, 2006. Thermal fluctuations of grafted microtubules provide evidence of a length-dependent persistence length. Proc. Natl. Acad. Sci. USA 103:10248–10253.

